# Tools for Drawing Informative Idiograms

**DOI:** 10.1101/2021.09.29.459870

**Authors:** Shoaeib Mahmoudi, Ghader Mirzaghaderi

## Abstract

Each species has a typical karyotype, which represents the phenotypic appearance of the somatic chromosomes including number, size, and morphology. Idiogram is a diagrammatic representation of the chromosomes showing their relative size, homologous groups and different cytogenetic landmarks. Chromosomal analysis of cytological preparations is an essential component of many investigations which involves the calculation of karyotypic parameters and generation of idiogram. Although various tools are available for karyotype analysis, here we demonstrate karyotype analysis using our recently developed tool named KaryoMeasure. KaryoMeasure is a semi-automated free and user-friendly karyotype analysis software that facilitates data collection from different digital images of the metaphase chromosome spreads and calculates a wide variety of chromosomal and karyotypic parameters along with the related standard errors. KaryoMeasure draws idiograms of both diploid and allopolyploid species into a vector-based SVG or PDF image file.

## 1. Introduction

In each species, *x* is the basic (monoploid) chromosome number, *n* is the gametic chromosome number and 2*n* is the zygotic or somatic chromosome number. For example, barley (*Hordeum vulgare*) has 2*n* = 2*x* = 14 chromosomes (RR genome) in each somatic cell, common wheat (*Triticum aestivum*) has 2*n* = 6*x* = 42 chromosomes constituting AABBDD genome and potato (*Solanum tuberosum*) has 2*n* = 4*x* = 48 chromosomes in each cell. In wheat and barley, the basic chromosome number *x* is seven while *x* in potato is 12. The coefficient of *x* shows ploidy level and 2*n* represents somatic or sporophytic (diploid multicellular stage of the life cycle of a plant or alga) chromosome number in a species. Hence, barley is a diploid (2*x*) with RR genome, common wheat is an allohexaploid (6*x*) with three AA, BB, and DD subgenomes and potato is an autotetraploid (4*x*) with AAAA genome.

Most species have monocentric chromosomes each with a localized centromere that divides the chromosome into small (S or p) and large (L or q) arms, even the centromere can be terminal. However, some species have holocentric chromosomes that lack localized centromere (Melters et al., 2012). In addition to the essential chromosomes, supernumerary B chromosomes (Bs) may additionally be observed in some species. Bs are not essential and their number varies between different individuals from 0, 1, 2, 3, or even more (Camacho et al., 2000; Marques et al., 2018).

In the case of allopolyploid plants, homoeologous relationships are also considered for karyotyping. Homoeologous chromosomes are chromosomes in the same species that originated by speciation and were brought back together in the same genome by allopolyploidization. Homoeologous chromosomes are commonly determined based on the previous studies of the meiotic behavior of the chromosomes or physical mapping of the homoeologous genes or sequences on the chromosomes.

Each species has a typical karyotype, which represents the phenotypic appearance of the somatic chromosomes including number, size, and morphology (Levitzky, 1931). Karyogram is the specific representation of homologous chromosome pairs from a metaphase chromosome picture where chromosomes are arranged in descending order. Idiogram is a diagrammatic representation of the chromosomes showing their relative size, homologous groups and varieties of chromosomal landmarks such as centromeres and banding patterns. Idiograms are used in many journal articles on a regular basis and in web interfaces to visualize varieties of genome and chromosomal information.

Karyotype analysis is an essential component of many cytogenetic investigations that involve measurement of karyotypic asymmetry indices and chromosomal characteristics such as number, size, position, and type of landmarks such as centromeres, secondary constrictions, and tandem DNA patterns (Heslop-Harrison and Schwarzacher, 2011). This information then may be used to distinguish homologous chromosomes and understanding the chromosomal and karyotypic evolution. By comparing the chromosomes of different taxa much can be learned about patterns and mechanisms of karyotype evolution and its significance for diversification and speciation (Weiss-Schneeweiss et al., 2008).

A lot of karyotype analyses have dealt with characteristics such as C- or N-banding patterns (Gill et al., 1991), *in situ* hybridization (ISH) banding patterns of repeated sequences (Lapitan *et al*. 1989; Albini and Schwarzacher 1992; Pedersen *et al*. 1996; Schrader *et al*. 1997), ISH of single-copy sequences (Shen *et al*. 1987; Peterson *et al*. 1999), or immunofluorescent localization of proteins on chromosomes (Anderson et al. 1999). The size and location of such landmarks are often quantifiable and are reflected in idiograms.

The traditional chromosomal measurement of different metaphase spreads and bringing the related data into tables is a slow and tedious process requiring intermediate photographic steps, manual statistical analysis, and manual drawing of the idiogram image. Interest in applying automation to the analysis of metaphase chromosomes dates back four decades and several computer applications have been developed so far to facilitate the accuracy of karyotype analyses (Armstrong et al., 1987; Fukui, 1986; Gilbert, 1966; Green et al., 1980; Green et al., 1984; James, 1983; König and Ebert, 1997; Oud et al., 1987). Some packages are also published for drawing the idiograms in R such as RIdeogram (Hao et al., 2020) and idiogramFISH (Roa and PC Telles, 2020). Currently, tolls such as MircoMeasure (Reeves, 2001), KaryoType (Altınordu et al., 2016), IdeoKar (Mirzaghaderi and Marzangi, 2015), ChromosomeJ plugin (Uhlmann et al., 2016) and, DRAWID (Kirov et al., 2017) are popularly used for the collection and analysis of cytogenetic data.

Here, we introduce our newly developed karyotyping tool called KaryoMeasure which is user-friendly and more flexible than our previously developed software: IdeoKar (Mirzaghaderi and Marzangi, 2015). KaryoMeasure is a software that facilitates chromosomal measurements and builds idiograms from digital images. KaryoMeasure is a semi-automated software and its use requires chromosome identification and numbering before karyotype analysis. In addition to extracting chromosomal and karyotypic parameters, KaryoMeasure is designed to additionally generate high-quality colored idiograms in vector-based PDF or SVG formats (Fig. 1).

**Fig. 1.**
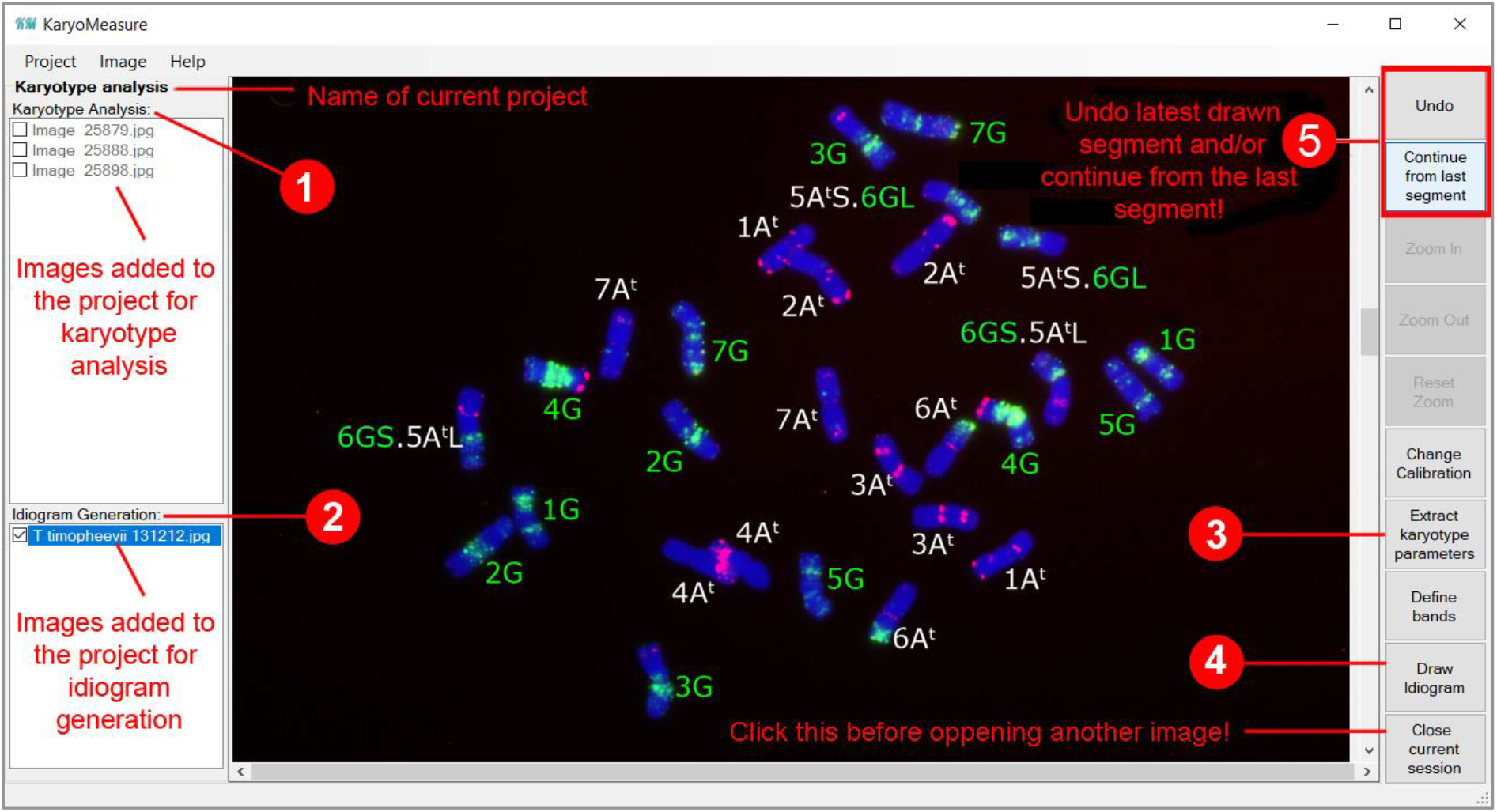
Using KaryoMeasure to trace extract karyotype parameters or generating idiograms. Images added to the project for karyotype analysis or Ideogram Generation are listed under Karyotype Analysis (**1**) and Ideogram Generation (**2**), respectively. After manual chromosome tracing, karyotypic parameters can be extracted (**3**) and idiograms can be generated (**4**). After reopening a project and selecting an image, it is possible to undo the latest drawn segment and/or continue from the latest point on the active image (**5**). If the latest point is a chromosome end, clicking on the ‘Continue from last segment’ key, reopens the end so that one can continue that chromosome. If you want to start a new chromosome, the previous chromosomes should be closed first because clicking on the ‘Continue from last segment’ key has already opened its end.

### Overall software design and architecture

KaryoMeasure software is used for the calculation of chromosomal and karyotypic parameters and to build idiograms. For this, a project is created at first, in which chromosome pictures are uploaded for karyotype analysis or idiogram generation via the ‘Image’ button in the menu bar. In this way, the pictures and upcoming activities on the pictures are automatically saved to the project so it is possible to continue analysis later after closing and reopening the project. Chromosomal and karyotypic parameters represented in Table 1, are extracted by KaryoMeasure. The nomenclature used for the description of type or morphology of monocentric chromosomes is that proposed by Levan *et al*. (1964) in which the chromosome type is defined based on the mean of arm ratio (Table 2). Stebbin’s asymmetry index is according to Table 3.

**Table 1.** Chromosomal parameters and karyotypic asymmetry indices along with their formula that are extracted by KaryoMeasure software.

| Chromosomal parameter or asymmetry index | Formula | Reference |
| --- | --- | --- |
| Chromosomal parameters |  |  |
| Arm ratio | $AR = L/S$ | |
| Chromosome length | $CL = L + S$ | |
| r-value | $S/L$ | |
| Relative length of chromosome | $RL\% = (CL/\sum CL) \times 100$ | |
| Centromeric index | $CI = S/CL$ | |
| Chromosome type | Defined Terms in Table 2 | (Levan et al., 1964) |
| Form percentage of chromosome | $F\% = (S/\sum CL) \times 100$ | |
| Karyotypic parameters or asymmetry indices |  |  |
| Total chromosome length of the haploid complement | $HCL = \sum CL$ | |
| Total form percentage | $TF\% = (\sum S / \sum CL) \times 100$ | (Huziwara, 1962) |
| Stebbins asymmetry index | A–C, 1-4 (Table 3) | (Stebbins, 1971) |
| Arano index of karyotype asymmetry | $AsK\% = (\sum L / \sum CL) \times 100$ | (Arano, 1963) |
| Intrachromosomal asymmetry index | $A_1 = 1 - [\sum_{i=1}^n (S_i / L_i) / n]$ | (Romero-Zarco, 1986) |
| Interchromosomal asymmetry index | $A_2 = s_{CL} / x_{CL}$ | (Romero-Zarco, 1986) |
| Symmetry index | $S\% = (CL_{min} / CL_{max}) \times 100$ | |
| Mean centromeric index | $x_{CI} = \sum CI / n$ | |
| Degree of karyotype asymmetry | $A = [\sum_{i=1}^n (L_i - S_i) / (L_i + S_i)] / n$ | (Watanabe et al., 1999) |
| Mean Centromeric Asymmetry | $x_{CA} = A \times 100$ | |
| Coefficient of variation of chromosome length | $CV_{CL} = (s_{CL} / x_{CL}) \times 100 = A_2 \times 100$ | (Paszko, 2006) |
| Coefficient of variation of centromeric index | $CV_{CI} = (s_{CI} / x_{CI}) \times 100$ | (Paszko, 2006) |
| Asymmetry Index | $AI = (CV_{CL} \times CV_{CI}) / 100$ | (Paszko, 2006) |

L: Length of long arm, S: Length of short arm, i: homologous group number, s: standard deviation, x: mean, n: haploid chromosome number of an individual or a taxon.

**Table 2.** Nomenclature of chromosomes based on Levan *et al*. 1964.

| Term | Centromeric position | Arm ratio | CI × 100 |
| --- | --- | --- | --- |
| M | Median point | 1 | 50 |
| m | Median region | 1-1.7 | 50-39.5 |
| sm | Submedian region | 1.7-3 | 39.5-25 |
| st | Subterminal region | 3-7 | 25-12.5 |
| t | Terminal region | 7-∞ | 12.5-0 |
| T | Terminal point | ∞ | 0 |

**Table 3.**
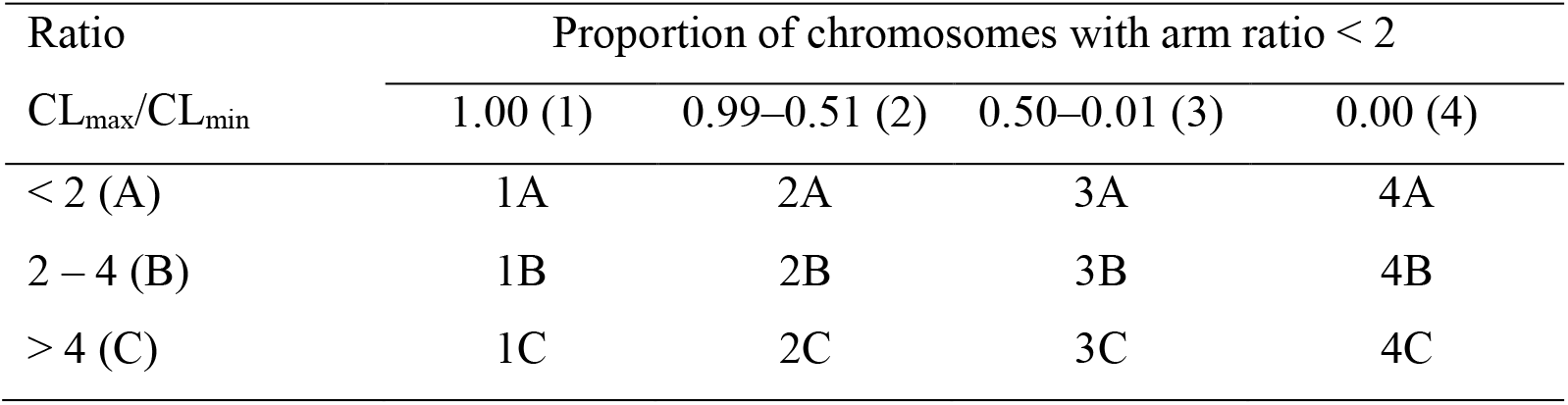
The classification of karyotypes in relation to their degree of asymmetry according to Stebbins (1971).

The chromosomal and karyotypic parameters are extracted after click-wise tracing of the chromosomes in KaryoMeasure. After that, all the parameters are automatically stored in a text file in the output folder of the project. The output text file includes karyotypic parameters, chromosomal parameters, and raw data which can be opened in Excel software as well (Fig. 2). KaryoMeasure calculates the mean and standard deviation of chromosomal parameters for all chromosomes of the same homologous group. These homologous chromosomes may be from one image or may belong to different images already opened in the project.

**Fig. 2.**
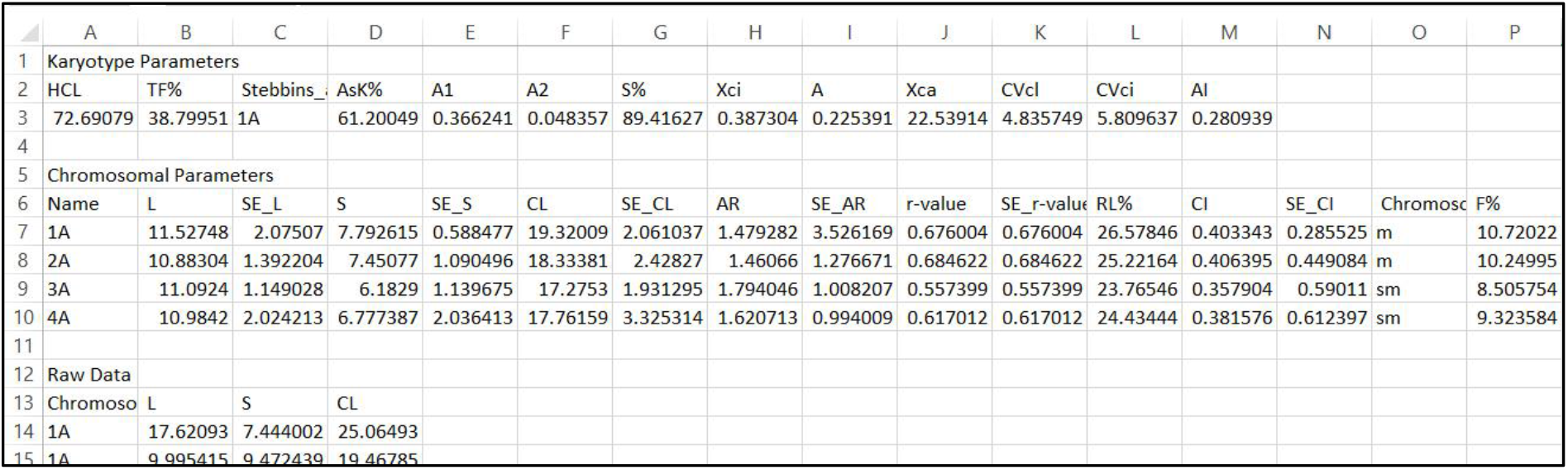
By adding images for karyotype analysis followed by manual tracing of chromosomes, a text file including karyotypic parameters, chromosomal parameters, and raw data will be added to the Output folder of the project.

## 2 Materials

### 2.1 Software and hardware requirements

1. KaryoMeasure software. KaryoMeasure is designed for standard personal computers with the operating systems Microsoft Windows 10. The latest version of the software can be downloaded from http://karyomeasure.uok.ac.ir/.
2. Mitotic metaphase chromosome pictures belonging to the same individual, same or different accessions or species depending on the aim of the study are required (*see* **Note 1**). KaryoMeasure supports input files with anyone of the jpg, tiff, bmp, and gif formats so there should be no need to convert the originally acquired images. It is possible to define a scale for all the opened images at once or a scale can be defined for each image separately, therefore it is not necessary that all the pictures be in the same magnification.
3. KaryoMeasure cannot identify homologous chromosomes automatically. Therefore, one needs to identify the homologs in advance. Homologs can be identified using a primary round of length measuring and/or by referring to arm ratios, banding patterns, and chromosome landmarks if available (*see* **Note 2**).

## 3 Methods

### 3.1 Karyotype analysis

1. **Defining a project:** When the software opened, a project can be created via the Project button of the menu bar and then clicking on New Project (Ctrl + N).
2. **Adding images:** For karyotype analysis, images can be imported via the ‘Image’ button of the menu bar and then the ‘Add image for Karyotype Analysis’. Different metaphase images can be added to the project as replicates for karyotype analysis. The imported images are arranged in the karyotype analysis field on the left side (Fig. 1). It is possible to remove any image along with the related data from a project via ‘Image’ and then clicking Remove Image’ button.
3. **Introducing scale:** Double click on one of the images and it will be opened. A scale needs to be defined by clicking on the start and end points of the scale bar of the image. A popup window appears asking for the actual length of the scale bar. Enter the corresponding size in µm and click on OK. It is possible to redefine the scale bar any time later via the ‘Change calibration’ key of the right sidebar.
4. **Assigning a name to each chromosome:** For tracing each chromosome, it is required to define the chromosome number and its genome (or subgenome) at first by right-clicking and pressing Start (or Ctrl + S). In the case of allopolyploid species, it is possible to assign the corresponding subgenome to each chromosome (*see* **Note 3**).
5. On each image, the user needs to trace the chromosomes using mouse clicking. The centromere and endpoint of the chromosome is defined via right-clicking or pressing Ctrl + C and Ctrl + E, respectively.
6. After tracing all chromosomes in an image, tracing can be continued in the other images, but the current session (e.g. the current image) must be closed before opening another image. The current session needs to be closed after tracing all chromosomes and before any further activity be conducted in the current project.
7. After tracing chromosomes in an image, ‘Close current session’ and double click on another image from the left pane to be opened. After tracing chromosomes in all images, karyotypic data can be stored in a text table file by pressing the ‘Extract karyotype parameters’ icon. The output text file has three parts: karyotype parameters, chromosomal parameters and raw data. Standard errors are also represented for each chromosomal parameter in the output table. The Notepad output file can also be opened in Excel software (Fig. 2).

### 3.2 Idiogram generation

1. **Adding images:** For idiogram generation, image(s) should be added to the project via the ‘Image’ button of the menu bar and then ‘Add image for idiogram generation’ (Fig. 1).
2. Different chromosome images can be added to the project for idiogram generation. The imported images are arranged in the ‘Idiogram Generation’ pane on the left side of the software (down area). Only one idiogram plate can be generated from each of these images.
3. Open one of the added images by double-clicking and introduce the image scale as mentioned in paragraph 3 of section 3.1.
4. If specific chromosomal landmarks such as centromere, C-, G-, N- or FISH bands, secondary constriction, 5S rDNA and 45S rDNA, etc. need to appear on the idiogram, a specific colour should be assigned to each of these landmarks via ‘Define bands’ icon before chromosome tracing (Fig. 3). Again, chromosome number and subgenome need to be defined for each chromosome by right-clicking and pressing Start (or Ctrl + S) before tracing each chromosome (*see* **Note 4**).
5. After tracing all chromosomes, clicking on the ‘Draw idiogram’ icon will display the idiogram control page where a number of adjustable controls are available for idiogram generation. On this page, the user can adjust chromosome height, chromosome width, centromere size and colour and scale bar size. It is also possible to sort the chromosomes by name or height in descending or ascending order. Usually, only one chromosome from each homologous pair is traced for idiogram generation (*see* **Note 5**).
6. In the case of species with holocentric chromosomes, no centromere is defined during chromosome tracing in KaryoMeasure. For telocentric chromosomes, the centromere and one chromosome end are overlapping. If tracing is started from the centromere end, the centromere is defined immediately after the start point (Fig. 5).
7. Another interesting option of ‘Idiogram generation’ is to use the karyotyping output table for producing idiogram. By selecting the karyotyping table in the idiogram control page, automatically, its idiogram can be produced (Fig. 6).

**Fig. 3.**
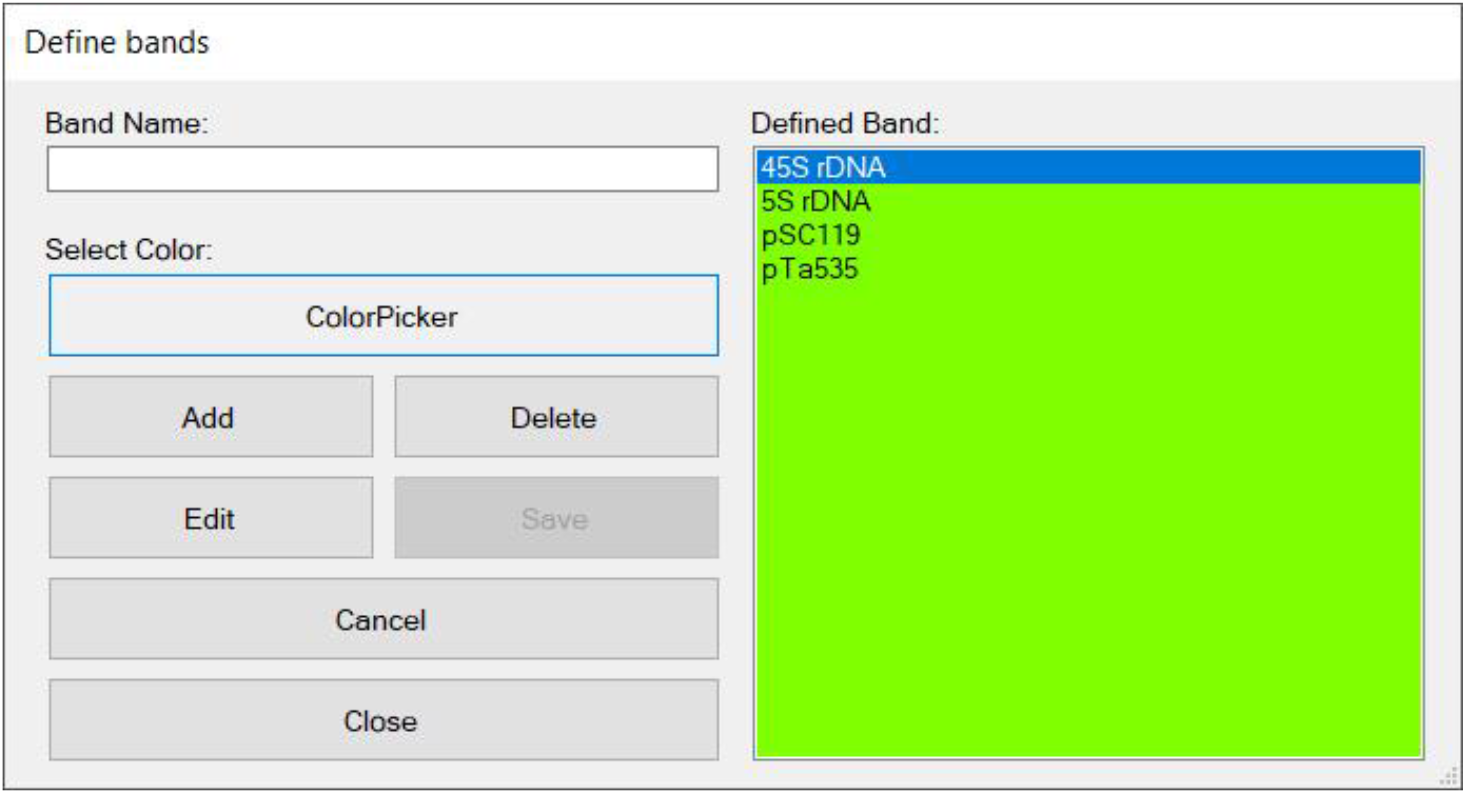
A color picker window will be appeared after clicking ‘Define bands’ icon where different colors can be assigned to different bands.

**Fig. 4.**
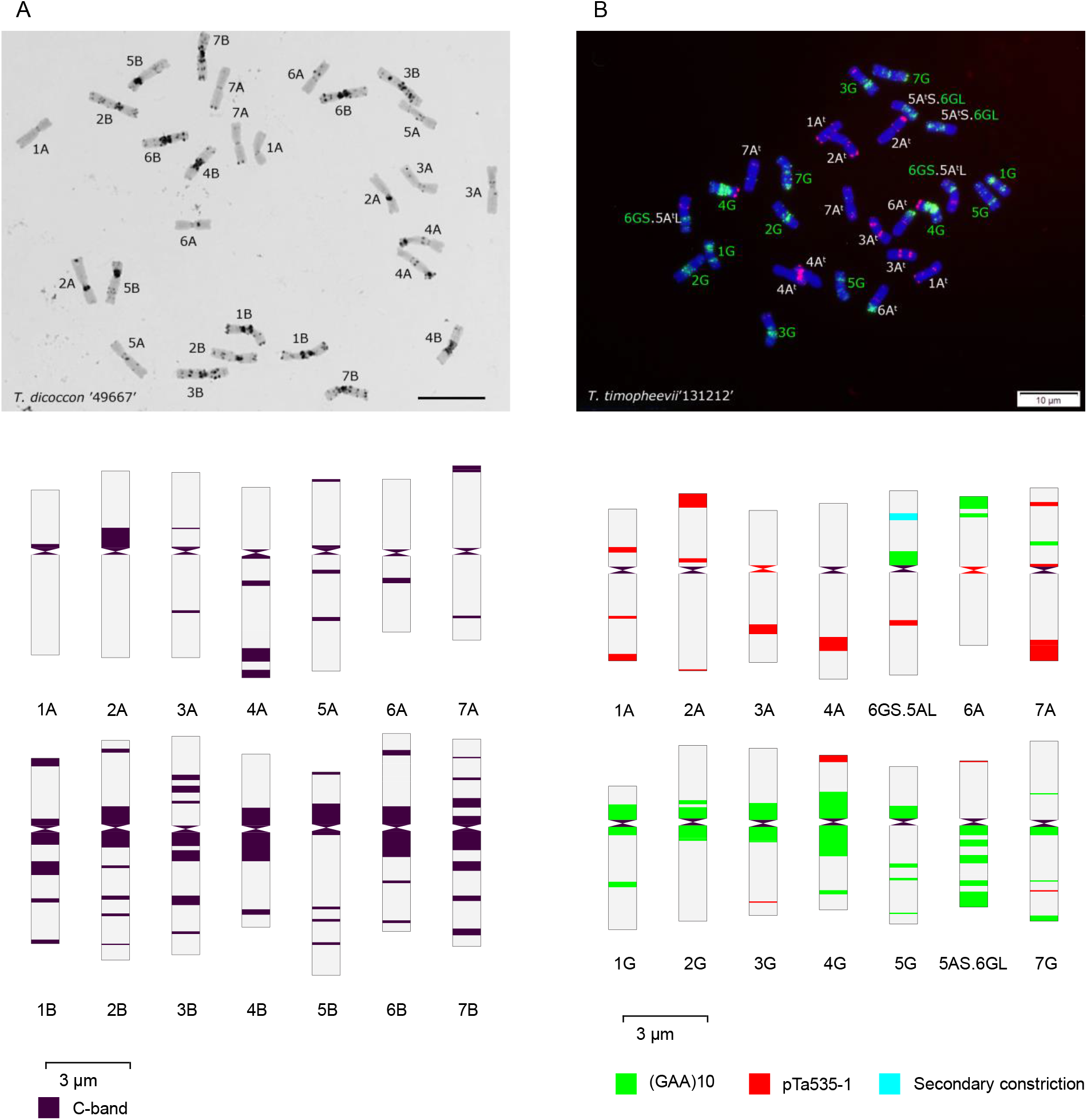
Generating idiogram of allopolyploid species using KaryoMeasure. C-banded idiogram of the *T. dicoccon* ‘49667’ (A) and FISH-banded idiogram of *T. timopheevii* ‘131212’ (B) generated by KaryoMeasure after tracing chromosomes. A scale bar is placed on the idiogram. From each homologous pair, only one chromosomes has been traced.

**Fig. 5.**
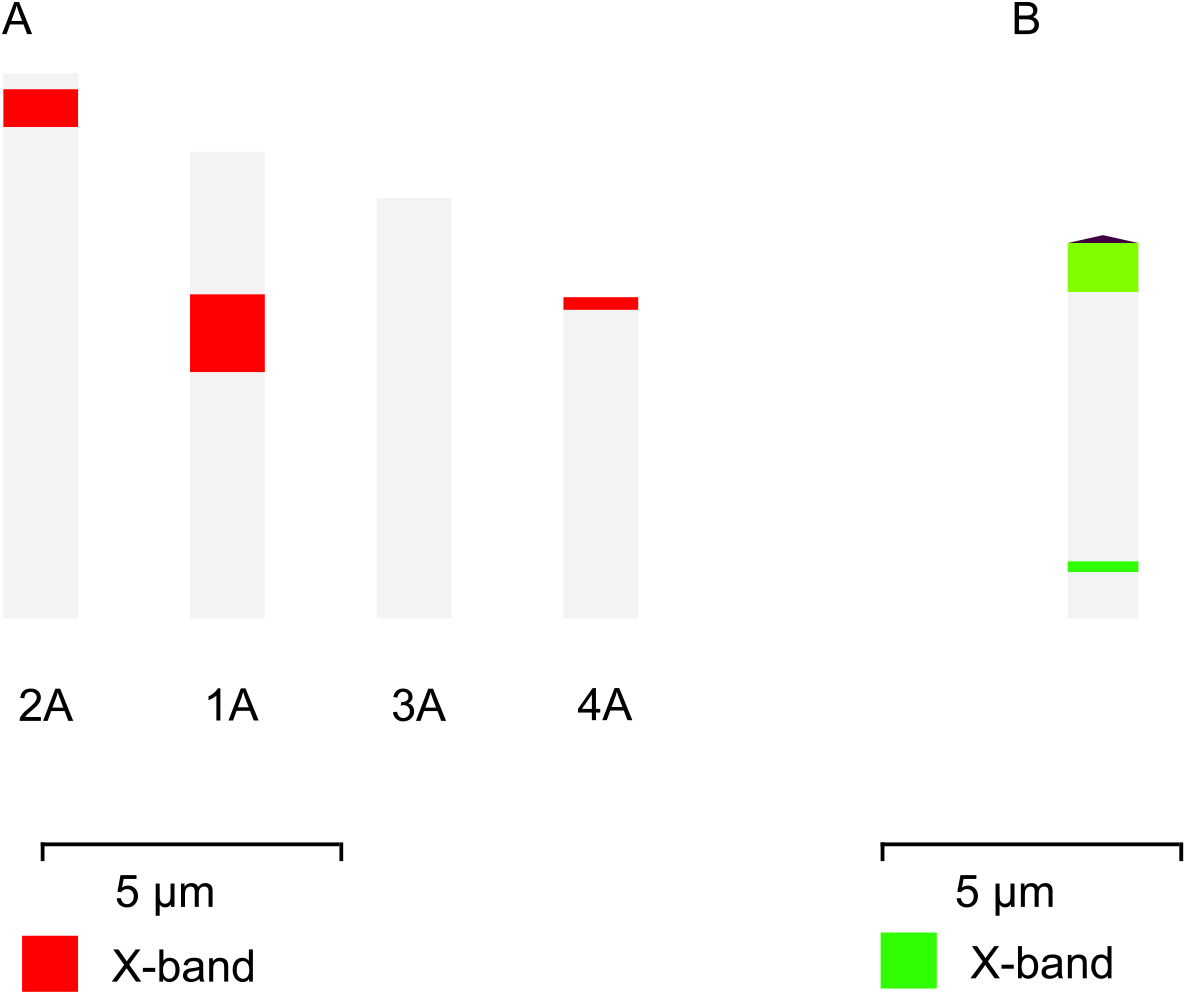
A) For the species with holocentric chromosomes, no centromere is defined during chromosome tracing in KaryoMeasure. B) For telocentric chromosomes, the centromere and one chromosome end are overlapping. If tracing is started from the centromere end, the centromere is defined immediately after the start point.

**Fig. 6.**
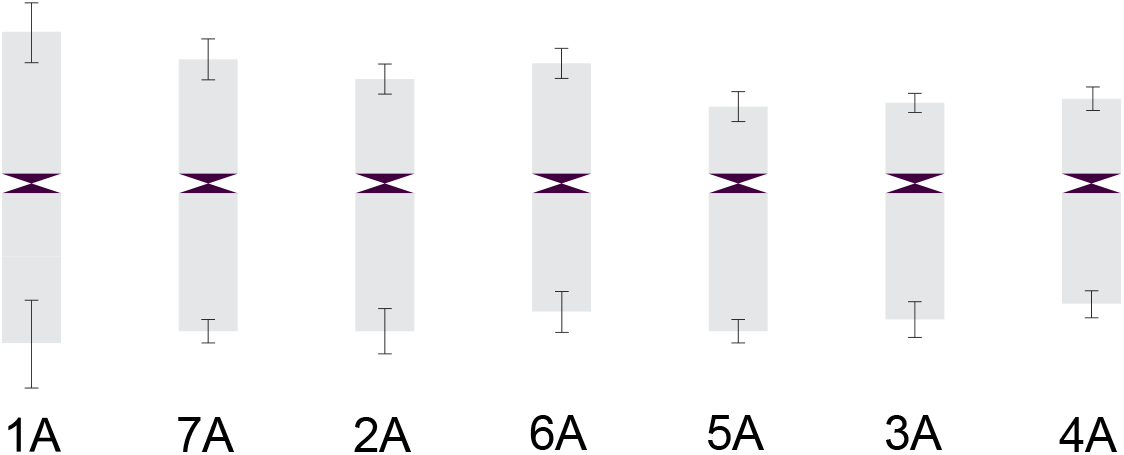
It is possible to use an output karyotype analysis table for idiogram generation in KaryoMeasure. An option is also available that adds standard errors of arms length to the chromosome endpoints.

## Notes

1. The whole or arm length of homologous chromosomes in the same or different cells can be very different because they are affected by a lot of factors such as pretreatment time and chromosome preparation method. Ideally, at least three high-quality metaphase spreads from three different cells or individuals should be analyzed for each sample.
2. Usually, it is not possible to identify all chromosomes only based on conventional staining techniques except the cases where chromosomes are morphologically enough heteromorphic. Looking for chromosomal landmarks such as chromosome sizes, arm ratios, secondary constrictions, size and position of heterochromatic knobs, and banding patterns resulting from Giemsa C-banding, N-banding, and FISH using repetitive sequences helps to identify all the chromosomes.
3. Allopolyploidy is very common among plant species. An allopolyploid has two or more different subgenomes in each cell. If the corresponding subgenome is assigned to each chromosome, chromosomes of each subgenome appear in a separate row in the output idiogram image.
4. If the species under study has B chromosomes, a letter different from that of normal chromosomes should be assigned to them. Consequently, Bs will be arranged under the normal chromosomes in the output idiogram picture (Fig. 7).
5. Producing SVG and pdf format is an advantage of KaryoMeasure because they are high quality and low weight and can further be processed in vector-based graphic softwares such as Inkscape or Adobe Illustrator. Editing colours and fonts and repositioning of objects are simply possible.

**Fig. 7.**
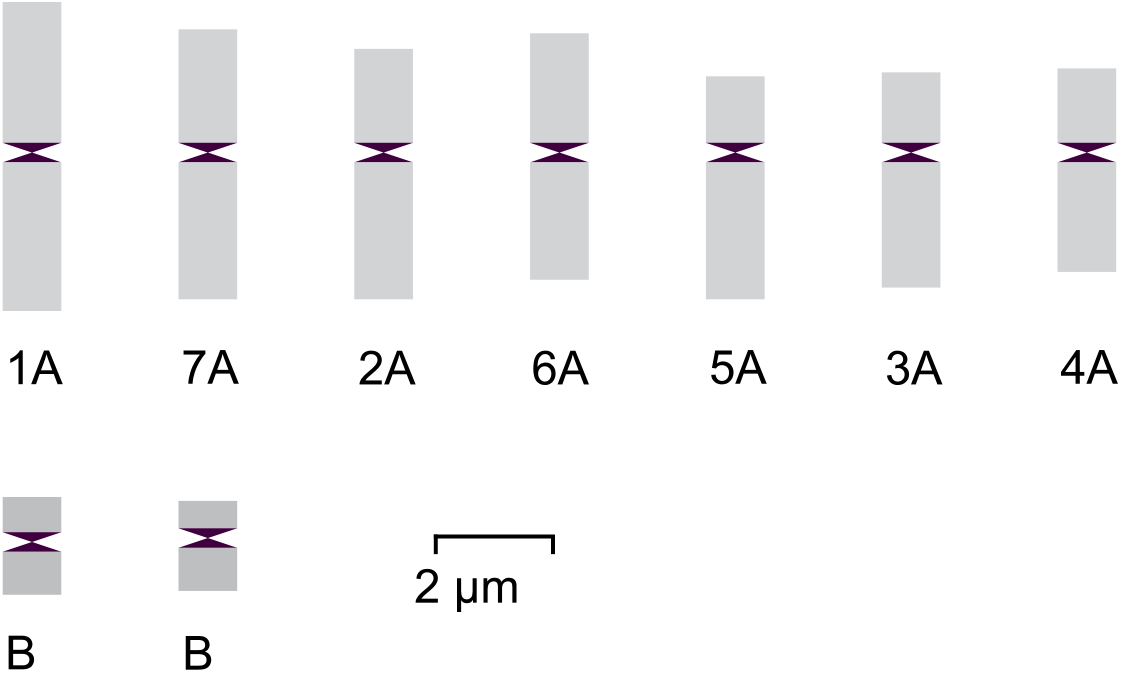
Idiogram of a species with 2 B chromosomes produced with KaryoMeasure and the Bs labels were edited in Inkscape (Note 6).

## Acknowledgement

This research was financially supported by the University of Kurdistan, Iran. The authors declare that they have no competing interests.

## Author’s contribution

SM developed the KaryoMeasure software. GM conceived and supervised the work and acquired chromosome images used in this study.

## Notes

http://karyomeasure.uok.ac.ir/

### Competing Interest Statement

The authors have declared no competing interest.

